# Genetic determinism and inheritance of differential DNA methylation profiles across three successive generations in quail

**DOI:** 10.64898/2026.09.16.751955

**Authors:** Stacy Rousse, Sophie Leroux, Rémi Séraphin, David Gourichon, Marta Gòdia, Ole Madsen, Sandrine Lagarrigue, Tatiana Zerjal, Frédérique Pitel, Sonia E. Eynard

## Abstract

Environmental exposures can induce epigenetic modifications that persist across generations, potentially contributing to the transmission of environmentally-induced phenotypes. However, the extent to which such persistence of molecular changes are maintained independently of genetic mechanisms remains unclear. In this study, we investigated the evolution and genetic determinism of DNA methylation (DNAm) variation across three successive generations of Japanese quails (G0 to G2) belonging to two epilines, defined according to whether their female ancestor had received genistein supplementation (epi+) or not (epi−). Using reduced representation bisulfite sequencing (RRBS), we characterised CpG methylation patterns and identified differentially methylated cytosines (DMCs) and regions (DMRs) between control and genistein-supplemented epilines. Out of 112,745 CpG sites analysed, no DMCs or DMRs were detected between epilines in the first generation following exposure (G0), whereas 621 DMCs (36 DMRs) were identified in G1, and 1,381 DMCs (101 DMRs) in G2. Despite this progressive increase of differential methylation sites, mean differences in methylation rate between epilines were globally small. Similarly, genome-wide genetic differentiation between epilines was very limited. Overall, SNP-based heritability of DNA methylation was 0.19 indicating that genetic variation accounted for only a small part of the global variation of DNAm levels. By contrast, DNAm at differentially methylated CpG sites was extensively under genetic control, with a mean SNP-based heritability > 0.5, and meQTL analysis further identified significant associations between SNPs and methylation levels. Together, these results indicate that local DNAm differentiation in quail is under substantial genetic regulation, while the limited genetic differentiation between groups suggests that genetic variation alone can not fully explain the progressive accumulation of methylation differences following an ancestral exposure to genistein. These results support DNAm as a plausible molecular candidate mechanism in the multigenerational response to environmental exposure and highlight the complex interplay between genetic and epigenetic regulations in the inheritance of phenotypes.

## Introduction

DNA methylation (DNAm) is an epigenetic mark participating in the regulation of gene expression. Extensively characterised throughout the tree of life, it involves the addition of a methyl group (CH_3_) to the DNA molecule. In vertebrates, this modification predominantly occurs at cytosines followed by guanines (CpG sites), which are arranged in a palindromic pattern (1–3). Mechanistically, *de novo* addition of the methyl group usually occurs at the fifth carbon of the pyrimidine ring of cytosines in CpG contexts and is catalysed by DNMT3A and B enzymes, while maintenance of the methylation states during DNA replication is catalysed by another enzyme of the same family, DNMT1 (4,5).

The methylome of vertebrates plays essential roles in the maintenance of cell integrity and functioning through its involvement in chromatin structure, DNA-protein interactions with transcription factors, and dynamic responses to environmental cues (5,6). In addition, the role of DNAm at CpG sites in the regulation of gene expression is context and tissue-dependent (7). For instance, methylation at CpG sites in proximity to transcription start sites (TSS) is generally associated with repression of gene expression, while methylation at CpG sites located within gene bodies can influence key functional properties, such as alternative splicing (3,8). DNAm is also determinant in maintaining tissue specificity as cells differentiate and commit to particular lineages (9,10).

Finally, without modifying the DNA sequence, DNA methylation is critical for normal development and has been extensively studied in a wide range of species as a potential mechanism linking genotype, environment and phenotype (11,12).

Changes in DNAm levels can be influenced by environmental factors, which can potentially contribute to variable phenotypic responses. Genome-wide differential methylation patterns have been observed in many distinct species in relation to environmental modification (13), including temperature, nutrition and husbandry practices. For example, the social differentiation of honey bees (*Apis mellifera*) into fertile queens or sterile workers is associated with DNAm differential levels. In fact, the sole difference of the feed they receive leads to changes in methylation patterns (14). More recently, Sourani and colleagues reported differences in the methylomes of adult chicken exposed to different breeding conditions during early life (15). Summer and spring dairy cows also express differential methylation profiles linked to the biological adaptation in response to heat stress (16). The importance of DNAm in contributing to phenotypic variability in response to environmental signals is therefore clearly established. However, most studies focus on longitudinal experiments, where the individuals directly exposed to the environmental stressor are the ones whose phenotypic responses are studied. In other words, the influence of DNAm as a vector of phenotypic variability is mostly described at a somatic level. For instance, multiple events of DNAm alterations following early-life stress have been associated with diseases in adulthood (9,17), highlighting the importance of DNAm in normal developmental, as well as its potential to retain signatures of past environmental exposures.

Such effects may also persist across generations. Several studies have reported the persistence of phenotypic modifications in the offspring of stressed animals, even when they are not themselves directly exposed to the stressor (4,18–20). Parental lifestyle, and particularly diet, has received considerable attention as a source of multigenerational effects. In pigs and mule ducks, dietary supplementation with methyl-donor compounds resulted in significant changes in body weight in later generations of descendants (21,22). Similarly, in beef cattle, methionine supplementation of mothers’ diet resulted in persistent methylation differences in calves, specifically in regions with genes involved in muscle development (23). Such examples collectively illustrate the relationship between temporary environmental exposure and phenotypic variation in subsequent generations and establish DNAm as a potential candidate mechanism to mediate environmental effects to the progeny (5). However, the precise transmission mechanism remains unclear in vertebrates. For example, following exposure of gestating F0 female rats to vinclozolin, an anti-androgenic endocrine-disrupting fungicide, subsequent analysis of the sperm of offspring revealed differentially methylated regions (DMR) between control and treated groups (24), even up to generation F13 (25).

Studying the multigenerational transmission of parentally-induced environmental effects holds different advantages and disadvantages depending on the focal parent. On the one hand, male germline is easily collected through sperm samples, it is abundant and the collection is non-invasive. On the other hand, even though evidence of non-genetic maternal effects have been evidenced (26), female germ cell collection remains difficult, their number is generally limited and the techniques are generally highly invasive.

Beyond these practical limitations, there are additional challenges in establishing that environmentally-induced DNAm changes are stably inherited and contribute to phenotypic variation. DNA methylation is, like other phenotypes, partly under genetic control (4,27–29). Consequently, the tight entanglement between genome, environment and epigenome complicates the study of epigenetic inheritance, as it requires distinguishing between the genetic inheritance of epigenetic states and epigenetic inheritance that occurs independently of DNA sequence. The integration of genomic and epigenomic information, combined with an experimental design that controls genetic resemblance between individuals is therefore essential to determine whether inherited phenotypic variation reflects classical genetic inheritance, environmentally-induced epigenetic regulation, or an interaction between both layers of biological information (4,19,30–34).

Livestock species are increasingly impacted by climate change, resource scarcity, immune challenges and changes in farming practices, making them good models to study how environmental factors may influence phenotypic variation through epigenetic modifications. The possibility of controlling the conditions in which livestock species are raised, together with the availability of detailed genetics, pedigree, and phenotypic information on traits ranging from growth to behaviour, make these species particularly suitable for studying genetic and epigenetic changes through time (35). Feed modification is amongst the easiest ways to alter an animal’s environment, as diets can easily be changed, adjusted or supplemented. Genistein, a component naturally present in soy, appears to be a relevant diet modifier for studying methylation changes, as it has been shown to alter natural methylation patterns in mice (36,37). In addition, genistein has been reported to affect growth and behaviour when administered at high doses (38,39).

The present study investigates the multigenerational genetic and epigenetic consequences of a single dietary genistein supplementation administered to female quail ancestors. The multigeneration design of this study encompasses both inter- and transgenerational effects, as quails were reared for three generations following the environmental exposure, such that the last generation was not directly exposed to the stressor, even in the germline (40). Previously, in Rousse et al. (2026) we identified phenotypic differences between the groups of genistein-supplemented individuals and controls, and detected an increasing contribution of the epiline effect to the variability of body weight across generations (41). A mirror-mating design was built to minimise the differences in genetic background of quails from both epilines and a simulation framework did not reveal any significant impact of genetic drift over three generations. To investigate whether the origin of the differences was a direct consequence of ancestral genistein ingestion, we examined molecular data from the same experimental population in the present study. Specifically, we collected red blood cells in 1,344 quails (from which 1,261 were analysed), as this tissue is easily accessible and can provide information on global systemic epigenetic variation thanks to the conservation of their nuclei in avian species (42). The evolution of DNA methylation patterns was studied across three generations in control (epi−) and genistein-supplemented (epi+) individuals to determine whether environmentally-induced methylation differences are transmitted across generations while accounting for potential changes in allele frequencies through genetic drift. The availability of molecular data for all individuals in all generations allowed the study of DNAm differential between epilines, inter- and transgenerational inheritance across groups. Large sample sizes for all generations also allowed to estimate genome-wide SNP-based heritability of DNAm. The size of the animal design also ensured reliable methylation quantitative trait locus (meQTL) analyses, enabling to assess the extent to which methylation levels at differentially methylated CpG sites are under genetic control.

## Materials and methods

### Experimental design

The quails included in this study originated from a multigenerational experimental mating design, in which two groups (thereafter called “epilines”) were maintained in parallel under standard rearing conditions following an initial diet disruption, as described in Rousse et al. (2026) (41). In brief, a mirror-mating design was established in which two full-sib females received the same standard diet with one of the two receiving additional genistein supplementation. This soy-derived compound served as the environmental perturbation applied to the population. Both females were mated with the same male. From this initial cross, two parallel epilines (control epi− and genistein-supplemented epi+) were reared for three generations, under standard breeding conditions (without any further changes in the environment), hence making the design suitable for a transgenerational study. This design aimed to maximise genetic resemblance between the two groups while preserving their respective inherited molecular characteristics. Blood samples were obtained from 1344 quails across 5 successive generations: G-2 founder generation (n=46); G-1 generation, in which half of the females received genistein supplementation (n=108); and the three subsequent generations G0 (n=404); G1 (n=374); and G2 (n=412). In these last three generations, each quail belonged to either the epi− or epi+ lines, according to whether their G-1 female ancestor had received genistein supplementation.

Animals were bred at the UE1295 PEAT (Nouzilly, France, https://doi.org/10.15454/1.5572326250887292E12) with official authorization for the animals (APAFIS#29977-2021010717073072 v5), and PEAT agreement (D371751).

### Reduced Representation Bisulfite Sequencing data

Methylation rates and genomic variations were obtained from Reduced Representation Bisulfite Sequencing (RRBS). Following the FAANG protocol (https://api.faang.org/files/protocols/samples/INRAE_SOP_blood_sampling_20210601.pdf), whole blood samples were centrifuged at 2,000g, for 5 min, at 4°C. The isolated red blood cells were then resuspended in a glycerol/PBS solution (80/20 v/v). DNA was extracted from red blood cells using a high-salt method (43). Libraries were then prepared using Hologic® Diagenode Premium RRBS Kit V2 (44) in accordance with the supplier’s instructions. Briefly, DNA was fragmented by digestion with the Msp1 restriction enzyme (5’ - CC/GG-3’ target sites). Following adapter and unique molecular identifiers (UMI) ligation, fragments were selected based on size. Samples with similar concentrations were then pooled together prior to bisulfite conversion. Next-Generation Sequencing was performed by Genewiz (Azenta Life Science, Leipzig) using NovaSeq2 to generate 150 bp paired-end reads. We developed a RRBS nextflow pipeline (https://forge.inrae.fr/genesis/rrbs_nextflow_pipeline) to process FASTQ files and obtain both genomic variants in VCF format and individual methylation rates in BED format. After quality control and adaptor trimming, alignment to the reference genome (<u>Coturnix japonica 2.1</u>, GCA_001577835.2) was performed using BISCUIT Version 1.7.1 (45). The software proceeds to a pre-treatment of the genome and creates three indexes based on the reference: Index1 transforms all cytosines to thymines, Index2 transforms all guanines to adenines and Index3 is the classical, unmodified, reference genome. This procedure serves for methylation calling and is adapted to bisulfite treatment. After alignment, reads were demultiplexed based on UMI sequences and only primary alignments with a mapping quality above 40 were retained. Samples with less than 100,000 mapped reads passing these quality thresholds were discarded.Joint methylation and variant calling across all remaining samples was subsequently performed using BISCUIT. CpG methylation sites and single nucleotide polymorphisms (SNPs) were then filtered separately.

### Methylation at CpG sites

After methylation calling, minimum read-depth filters were applied to each sample individually. CpG sites with a minimum coverage of 10 uniquely mapped reads were retained. A maximum threshold was also applied to each sample, to discard CpG sites with a read depth exceeding the 99.9th quantile of CpG depth distribution. Samples with a CpG site missing rate over 90% were discarded. Lastly, only CpG sites shared by 80% of samples were kept for downstream analyses. A combination of two lists of SNPs was used to filter out CpG sites that overlap known SNP positions, as bisulfite conversion can be confused with a SNP C > T. These two lists comprised, on the one hand, the SNPs identified by the pipeline described above and, on the other hand, SNP positions identified from an Oxford Nanopore sequencing experiment performed on 23 quails from the same lineage (46). Finally, CpG sites with a DNA methylation rate standard deviation (s.d) ≥ 5%, therefore informative for differential analysis, were retained (see Supplementary Figure1 for details on the filters applied). CpG sites were analysed when located on autosomes (chromosomes 1 to 28 and linkage groups ChrLGE22C19W28_E50C23 and ChrLGE64), the sexual chromosome (Z), the mitochondrial genome (MT) as well as unannotated scaffolds.

### Genotypes

SNPs obtained by variant calling from RRBS data were first filtered in each sample for a minimal quality of 10 and a minimal read depth of 5. A maximum depth threshold was also set at the 99.9th quantile of SNP depth distribution. After concatenating filtered data across all individuals, positions detected in fewer than 90% of samples were discarded. Only SNPs with a minor allele frequency (MAF) > 0.01 were then kept. SNPs were analysed when located on autosomes (chromosomes 1 to 28 and linkage groups ChrLGE22C19W28_E50C23 and ChrLGE64) as well as unannotated scaffolds grouped in a synthetic unknown chromosome.

### Genomic features

The definition of genomic features was done following the annotation available on Ensembl (release 116, GCA_001577835.1) for *Coturnix japonica*. The affiliation of each CpG site to a genomic feature was performed using the R package GenomeFeatures (https://forge.inrae.fr/aurelien.brionne/GenomeFeatures). Genomic feature association was performed for all the CpG sites annotated on the quail genome and for the CpG sites present in our RRBS analysis. Relative enrichment of RRBS CpG sites across genomic features was then tested by comparing the whole-genome and RRBS CpG site distributions using odds ratio test.

### Differential Analysis

Differentially methylated cytosines (DMC) and differentially methylated regions (DMR) were identified using the R package Dispersion Shrinkage for Sequencing data (DSS version 2.54.0) (47) in R 4.4.0. Notably, not all individuals displayed at least 10 reads across the 112,745 CpG sites kept after all above-mentioned filters. DSS relies on a beta binomial model to take the sequencing depth into account together with the number of methylated reads. Therefore, this software allowed us to retrieve the entirety of methylation information for those CpG sites. In fact, the posterior distribution and statistical tests are computed taking into account the poor amount of information brought by some observations. We fitted a multifactorial model per generation including the effect of sex, epiline, batch and the sex-epiline interaction. In brief, DSS multifactor model uses an ‘arcsine’ link function to link expected methylation rates (beta distribution mean) and a linear combination of the covariates. Differentially methylated regions (DMR) were subsequently identified for epiline. Both DMC and DMR were considered significant at a false discovery rate (FDR) threshold equal or below 0.05. For this analysis, only generations post-genistein ingestion (G0 to G2) were considered. The sample sizes in each epiline per generation were for G0: n_epi−_ = 159 and n_epi+_ = 202; for G1: n_epi−_ = 161 and n_epi+_ = 195; for G2: n_epi−_ = 186 and n_epi+_ = 213.

The validity of DMC detection was verified using Monte Carlo permutation tests. The Monte-Carlo permutation p-value (probability to detect as many DMC by chance if groups had no real impact) was computed as 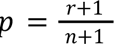 where *n* is the total number of simulations and *r* is the number of permutations for which we detect a similar or higher number of DMCs than with the original dataset (without permutation) (48,49).

### Functional annotation and enrichment analysis

To obtain a complete inventory of genes overlapping DMC/DMR, positions were first overlapped with gene positions from Ensembl 116 using the bedtools intersect option (50). Thereafter, genes were annotated using the R package biomartr (51) for cross-species mapping (quail, chicken, and human loci). Functional annotation was performed using UniProt (52), on the subset of these genes with a recognised human orthologue, to support gene-level functional interpretation. A functional enrichment using g:Profiler (53) was also performed.

### Fixation index (F_ST_) between epilines across generations

The changes in allele frequencies across three generations were studied by comparing the bi-allelic diversity between epi− and epi+ (n = 1,180) using Weir and Cockerham estimators of F_ST_ (54) with Plink software (version 2.00a4), in each generation. F_ST_ was also computed between G0 and G2 (n = 806) to check the overall allelic differences accumulated over the total time span of the experiment after genistein ingestion.

### Estimation of DNA methylation heritability

SNP-based heritability of the methylation at CpG sites was estimated using the restricted maximum likelihood (REML) option implemented in Genome-wide Complex Trait Analysis (GCTA) software, version 1.94.1 (55). The variance components were estimated in a linear mixed model:

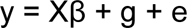

where y represents the vector of DNAm at a given CpG site, Xβ represents fixed effects included in the model (epiline, sex, generation and batch), g represents additive genetic effects with covariance structure defined by a genomic relationship matrix (G) computed across all the available individuals (from G-2 to G2; n = 1,334 after filters on read alignment and quality), and e represents residual effects. The additive genetic and residual variances were estimated as g∼N(0,Gσ_g_^2^) and e∼N(0,Iσ_e_^2^), respectively, where σ_g_^2^ is the additive genetic variance, σ_e_^2^ is the residual variance and I is an identity matrix. SNP-based heritability was subsequently calculated as 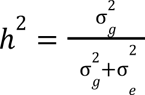. The genomic relationship matrix (G) was also built in GCTA, using the 38,897 SNPs located on autosomes, and added in the model to account for similarity between individuals, based on their SNP genotypes. Sequenced individuals from all five generations were used at this step (N=1,334).

### Methylation Quantitative Trait Loci (meQTL) analysis

Associations between variation in DNAm levels and SNP genotypes were analysed using the mixed linear model association (MLMA) implemented in GCTA. The association study was only conducted at CpG sites of interest, identified in the methylation differential analysis. For each SNP, the mixed linear model fitted was:

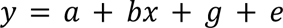

where *y* is the vector of DNAm levels at an identified CpG site across individuals, a is a mean term, *bx* represents the fixed effect of the tested SNP, *g* accounts for the random polygenic effect of all other SNPs captured by the GRM, and *e* represents the residual error. The null hypothesis that the tested SNP had no association with the phenotype was evaluated after accounting for multiple testing. The thresholds for significance were determined differently according to the type of association. For the detection of cis-association, the threshold 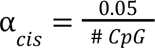 was applied for all SNPs located on the same chromosome as the CpG considered, within a 1 Mb window. For other SNPs, located outside the 1Mb window on the same chromosome, or on a different chromosome, the threshold 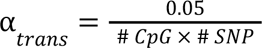 detection of trans-association.

## Results

### CpG sites and DNA methylation rate distribution

On average, 14,8 M reads were obtained per sample, across 1,344 individuals sequenced. Bam files contained an average of 4,138,656 uniquely mapped paired-end reads after quality control for downstream analyses. The total number of CpG sites detected with RRBS was 1,608,643. After the filtering steps applied in the pipeline, 481,496 CpG sites were kept (Supplementary Figure1). By removing SNPs, this number was reduced to 457,724. The additional filtering step to keep CpG sites showing variability (standard deviation ≥ 5%) led to the conservation of 112,745 CpG sites for analysis. This final dataset represented 7.01% of the total number, across 1,261 quails included in the experimental design. In our dataset, keeping CpG sites showing variability led to discarding many CpGs with extreme DNAm rate, as well as sites displaying little to no difference in average DNAm rate between epilines (Figure 1A).

**Figure 1:**
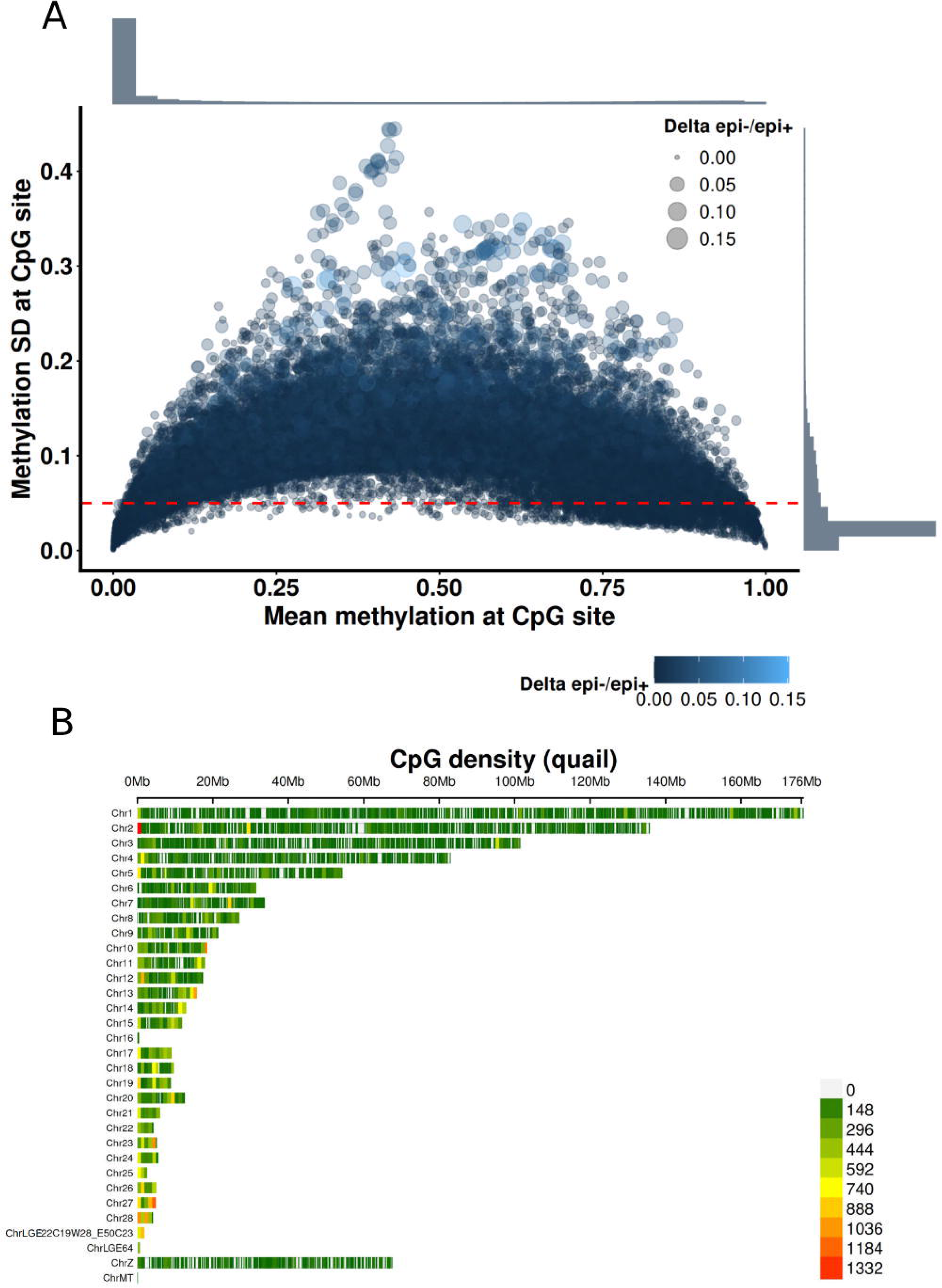
Mean methylation rate as a function of methylation standard deviation and CpG sites distribution from RRBS. A: Scatter plot of CpG methylation rate, showing the mean methylation rate (x-axis) as a function of the standard deviation (s.d.) (y-axis) for all CpGs passing the standard filters of the RRBS nextflow pipeline and SNP overlap filtering (n = 457,724). CpG sites above the red line (s.d. ≥ 0.05), were kept for downstream analyses (n =112,745). The upper and right-sided histograms represent the overall distribution of methylation rates and standard deviations, highlighting a strong proportion of low methylation levels, with little variability. The colour and size of dots represent the mean methylation difference between epi− and epi+ individuals per CpG site. B: Density map showing the distribution of CpG sites along the autosomes across bin sizes of 1 Mb (chromosomes 1 to 28 and linkage groups ChrLGE22C19W28_E50C23 and ChrLGE64), the sexual chromosome (Z) and the mitochondrial genome (MT).

The median depth was 32.5 reads per CpG site. Overall, the analysed CpG sites were distributed across most chromosomes, with higher densities at chromosome extremities. Microchromosomes displayed a higher concentration of CpG sites compared to macrochromosomes (Figure 1B). Although the analysed sites represented only a small proportion of CpG sites in the quail genome, they were well-distributed across the genome.

### Genomic features and enrichment analysis

DNAm rates varied across genomic features on autosomes, with exons, coding sequences (CDS) and 3’UTR regions exhibiting an elevated proportion of highly methylated sites relative to sites with low methylation rates. Promoters, introns, 5’UTR, and intergenic regions displayed the opposite pattern (Figure 2). In general, epi+ and epi− did not show specific DNA methylation distribution patterns across genomic features and globally displayed similar profiles.

**Figure 2:**
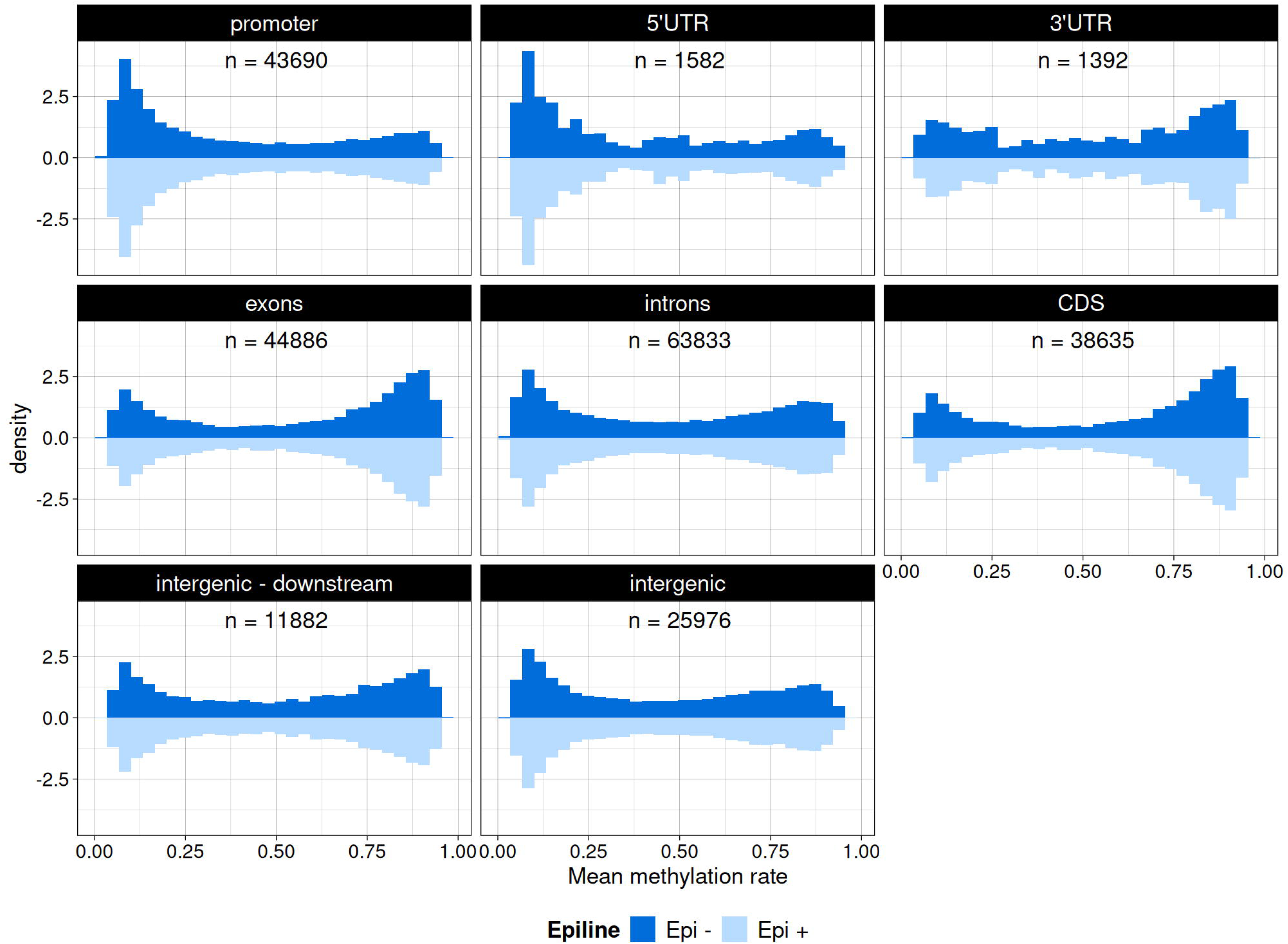
Distribution of mean methylation rate per genomic features on autosomes. Upper histograms represent individuals of the epi− group while bottom histograms represent epi+. N values in each grid displays the number of CpG sites annotated for the concerned genomic feature, knowing that one CpG site can be considered in several features (e.g. A CpG site can be found in the promoter of one gene and in the body of another gene).

Genomic regions enrichment was assessed by comparing the annotation of total CpG sites on autosomes on the quail genome with RRBS generated CpG sites used in our analyses. While about half of all CpG sites in the quail genome were located within introns, less than 30% the CpGs analysed by RRBS fell into intronic regions, with a corresponding enrichment in regulatory regions (promoters, 5’UTR, and 3’UTR), coding regions (CDS and, exons), as well as intergenic regions located up to 1000 bp downstream of coding sequences (Figure 3A). In addition, a significant enrichment of these regions in the RRBS dataset was detected, with odds ratios ranging from 1.4 for 5’UTRs to 2.9 for CDSs. Conversely, intronic and intergenic regions were under-represented, with odds ratios of 0.65 and 0.61, respectively (Figure 3B).

**Figure 3:**
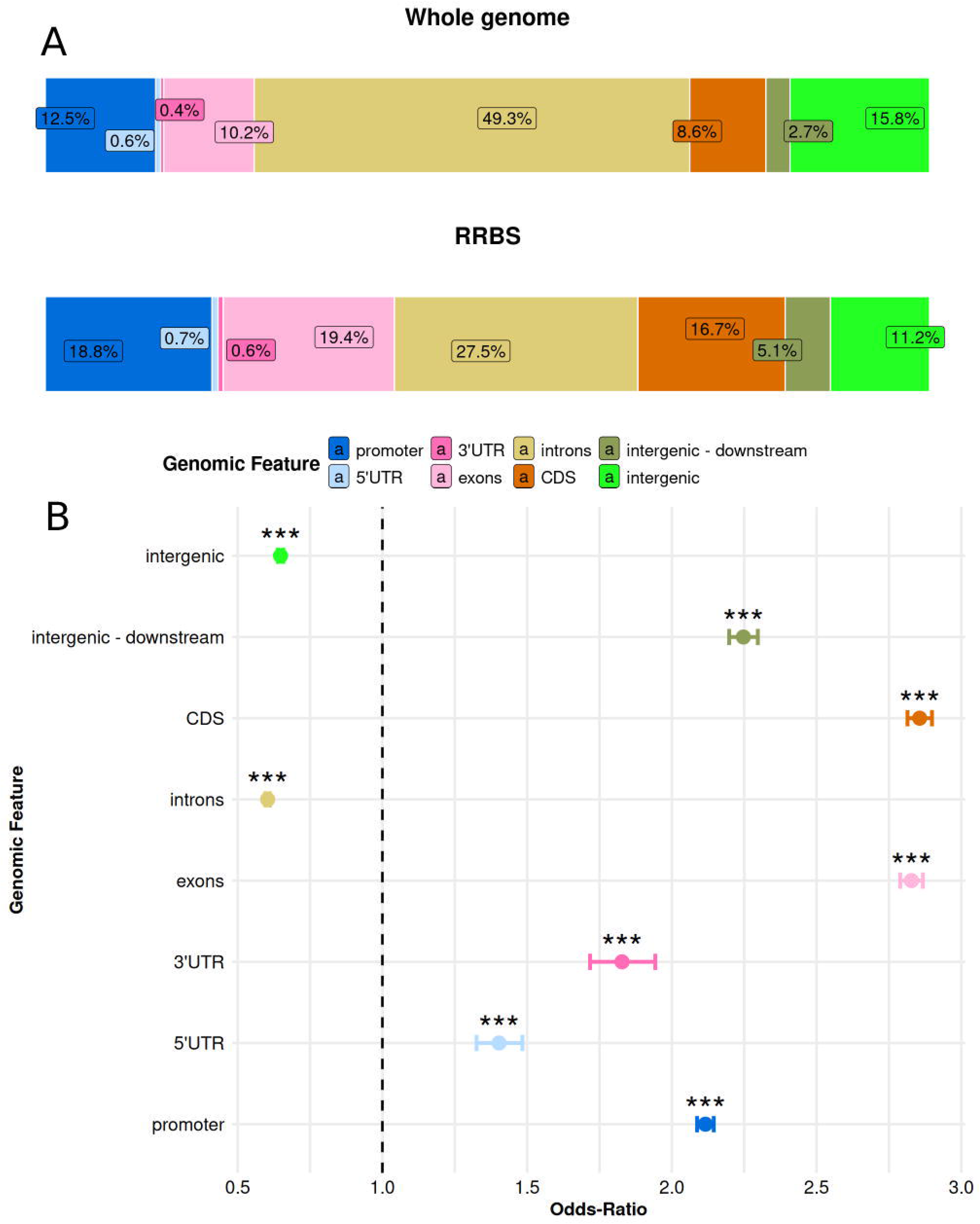
Genomic features enrichment for CpG sites obtained with RRBS compared to CpG sites across the whole quail genome. A: CpG site repartition across genomic features for the whole-genome (upper-strip) and for RRBS post filters (bottom-strip). B: Odds ratio test results for relative enrichment in genomic features of RRBS compared to whole-genome distribution of CpG sites.

### Methylation differential analysis

The inclusion of the sex-epiline interaction did not reveal any Differentially Methylated Cytosine (DMC), regardless of the generation. This interaction term was therefore removed from the model. After this adjustment, a total of 2002 DMCs associated with the epiline effect were identified: none in the first generation following genistein ingestion (G0), 621 in G1 and 1381 in G2. Among these 166 DMCs were common to G1 and G2 (133 on known chromosomes, 33 on unknown scaffolds). Because of the large sample sizes in the generations studied, the differential analysis revealed significant DMCs even at CpG sites showing very small differences in mean methylation levels between the groups (< 1%; Figure 4). Among the 166 common DMCs identified in G1 and G2, the majority of sites that were hypomethylated in G1 were also hypomethylated in G2, suggesting conservation of the methylation patterns between these two generations. The same was observed for hypermethylated sites (Table 1; Supplementary Figure2).

**Figure 4:**
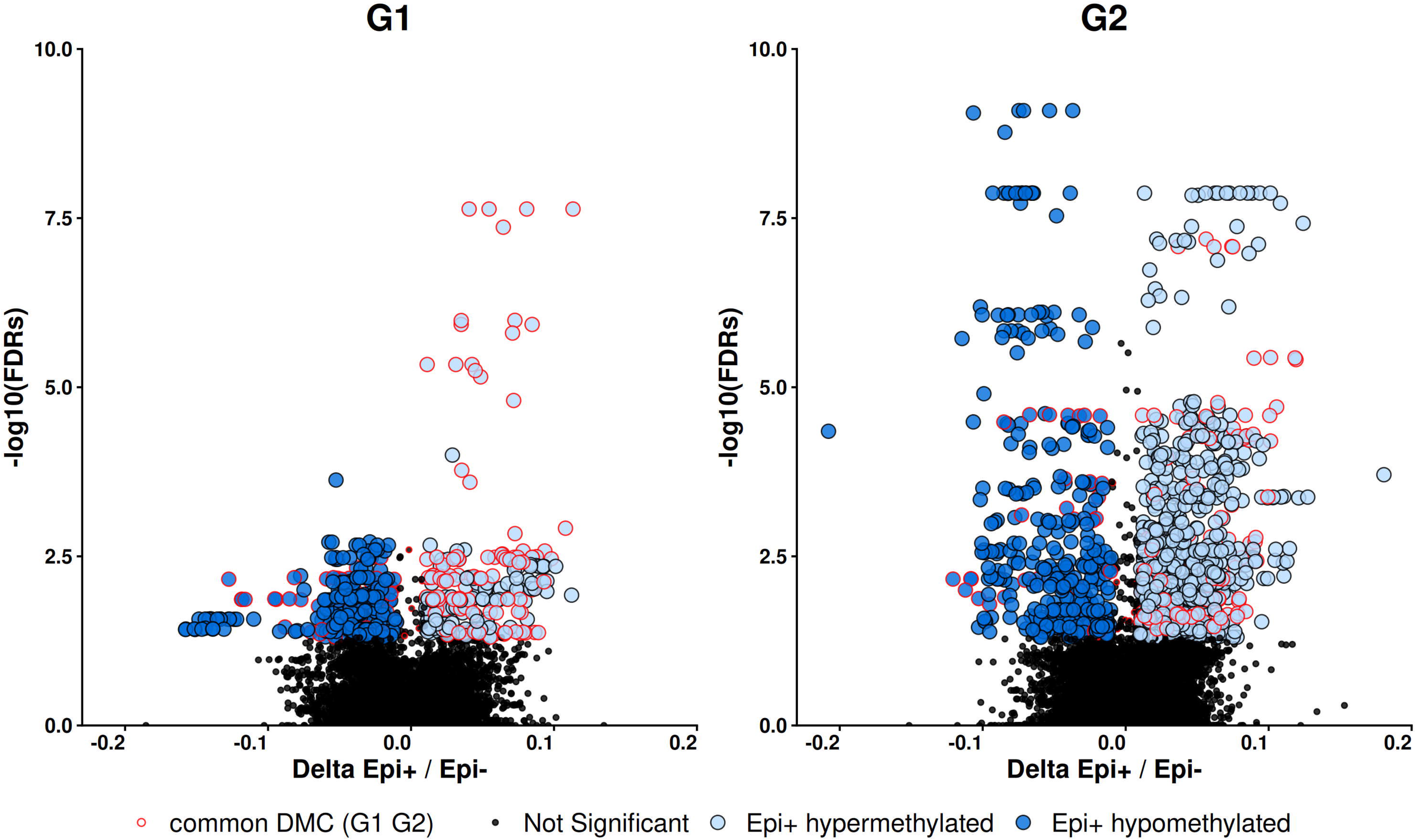
Differential Methylation between epilines. A: Volcano plot of statistical significance (-log10(FDR) versus the magnitude of the mean methylation difference between epilines (Delta Epi+/Epi−) in G1 (left graph) and G2 (right graph). DMCs that were associated with a Delta ≥ |0.01| (1% mean methylation difference between epi+ and epi−) are highlighted in dark blue (hypomethylation of Epi+) or in light blue (hypermethylation of Epi+). The CpGs with a red outline correspond to DMCs common between G1 and G2.

**Table 1.**
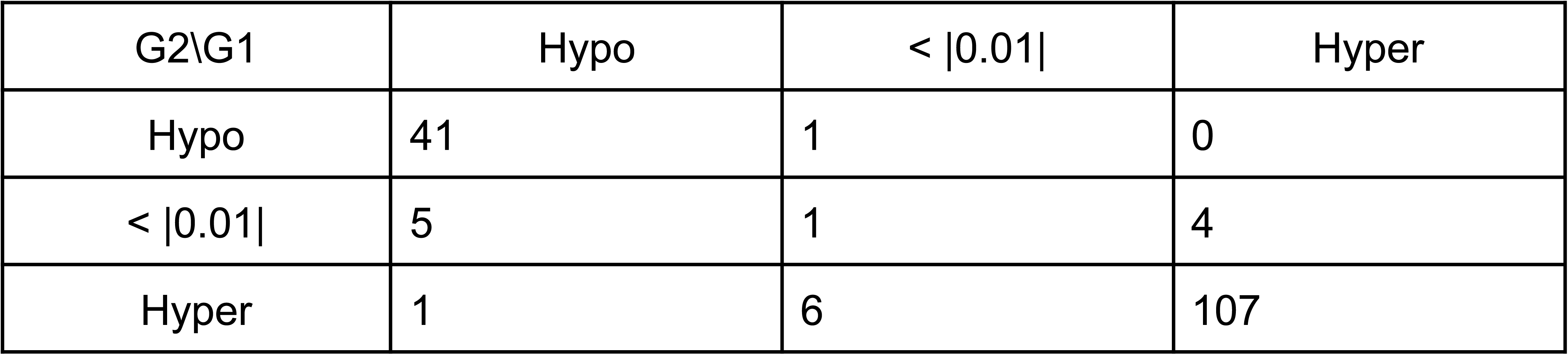
Methylation differential patterns between epi+ and epi− in common DMCs between G1 and G2. *Rows: G2, Columns: G1*.

Differentially Methylated Regions (DMRs) were also detected in G1 and G2 (36 and 101, respectively). The overlap between these DMRs revealed 13 common regions. Generally, all DMCs were contained within DMR regions (Supplementary Table1).

For the remaining factors included in the model, a substantial number of DMC associated with sex were identified in all generations, and these CpG sites were largely located, as expected, on chromosome Z (see Supplementary Figure3). In all three generations, a large number of DMC associated with the batch effect were also detected. This result confirms the importance of including batch in the model to account for variation unrelated to biological effects of interest. The batch differences may arise from technical variability during sample processing, library preparation or sequencing.

Permutation testing confirmed that the observed DMCs were not a result of a stochastic phenomenon, despite their limited number. The number of observed DMCs for each factor appears significantly higher than that obtained from simulations using randomly permuted metadata supporting the significance of the found DMCs (see Supplementary Figure4).

### Annotated genes associated with DMCs/DMRs

The genes associated with differential methylated cytosines and differential methylated regions are listed in Supplementary Tables2-5. The number of annotated genes increased from generation 1 to generation 2 for both gene bodies and promoters, consistent with the larger number of DMCs and DMRs identified in G2 compared to G1.

Gene Ontology (Biological Process) enrichment analysis was performed separately for each list defined according to DMC/DMR status, generation (G1, G2 or common to both generations), and genomic location (gene body or promoter), using genes covered by RRBS sequencing as the background. No significant enrichment for any functional category was identified after correction for multiple-testing.

The intersection of common DMC and common DMR lists narrows the candidate set to a small group of loci differentially methylated across both generations, at both single-cytosine and regional resolution: four genes in the gene body (*DHPS*, *GCDH*, *JPH3*, *WDR83*) and four genes in the promoter (*ARHGDIG*, *HYAL3*, *NAA80*, *NUDT16L1*).

### Fixation index (F_ST_) between epilines across generations

F_ST_ was computed to assess genetic differentiation between different quail subsets. First, all individuals of the G0 were compared with all individuals of G2. The distribution of F_ST_ along chromosomes revealed genetic differentiation between quails from G0 and quails from G2 at specific genomic locations, particularly at the beginning of chromosome 2, and on chromosomes 4 and 15 (Figure 5). Similar patterns were observed when G0 and G2 were compared separately within each epiline, suggesting that these allelic differences accumulated across generations rather than being specific to one epiline (Supplementary Figure5). However, F_ST_ values in these regions were moderate, always below 0.2. By contrast, no apparent signature of genetic differentiation between epilines was revealed. The overall values of F_ST_ between the two epilines remained lower than 0.1 genome-wide. Mean F_ST_ values F_ST_ in both subsets were 0.003 and 0.004 for G0/G2 and epi−/epi+, respectively (Supplementary Table6).

**Figure 5:**
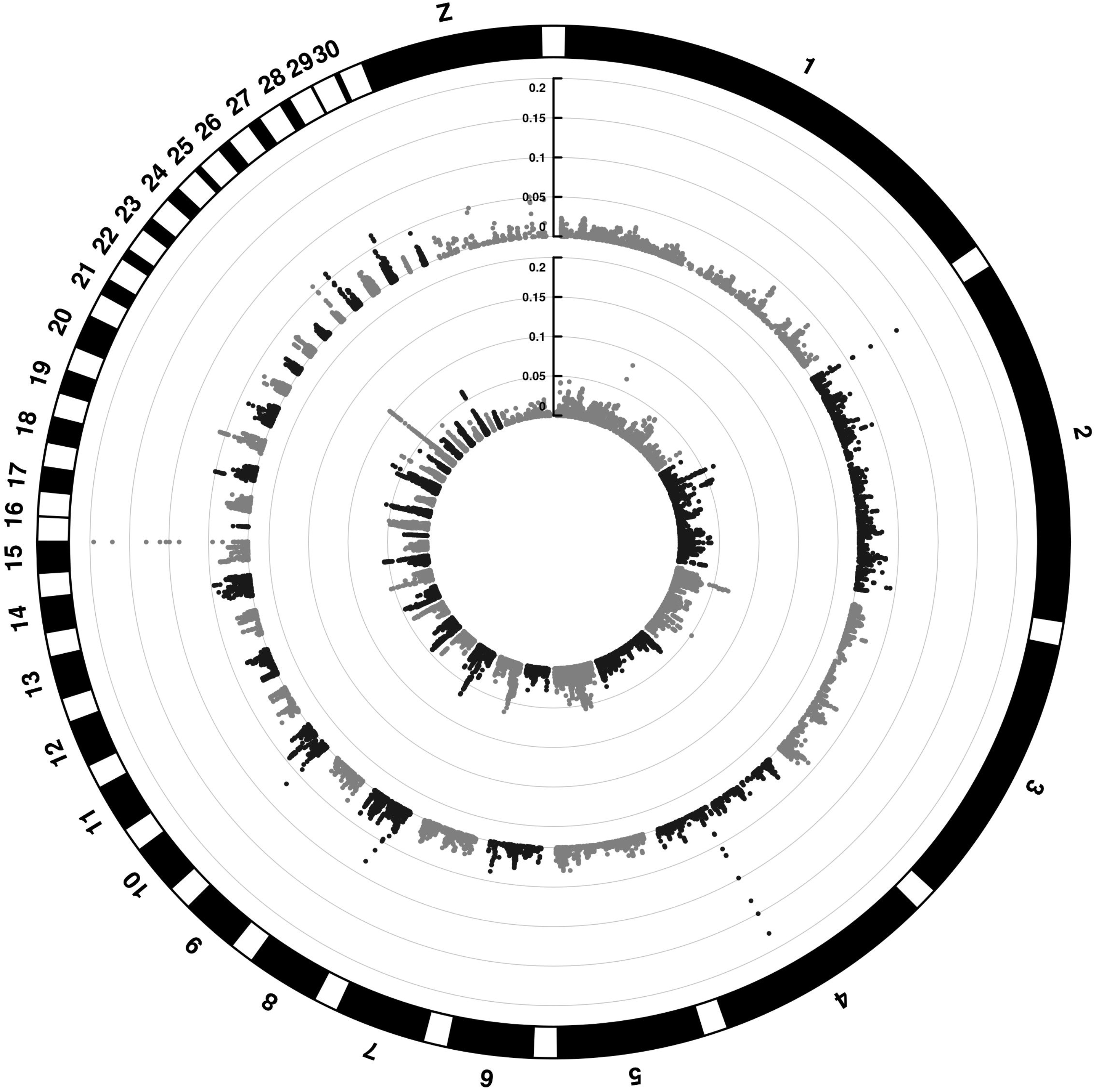
Genome-wide F_ST_ distribution fromRRBS-called SNPs. F_ST_ values along the quail genome reflecting allele frequency differences between the subsets compared. On the inner track: epi− individuals versus epi+ individuals (G0 and G2 confounded). On the out track: G0 individuals versus G2 individuals, regardless of their ancestral exposure to genistein. The y axis portrays the F_ST_ value. The distribution in all autosomes is represented (chr1-chr28), chromosomes 29 and 30 correspond to the linkage groups chrLGE22C19W28_E50C23 and chrLGE64 respectively.

### DNA methylation heritability estimates

The SNP-based heritability of DNA methylation levels at CpG sites in quail red blood cells was estimated at 0.19 on average (Figure 6A). While the majority of CpG sites showed little to no heritability, a few sites displayed high heritability estimates, reaching values close to one. Among these sites, CpG sites that were identified as DMCs in G1 or G2, showed higher heritability estimates than the overall set of CpGs sites, with average values of 0.57 and 0.54, respectively. DMCs common to both G1 and G2 showed an even higher average heritability of 0.63 (Figure 6B).

**Figure 6:**
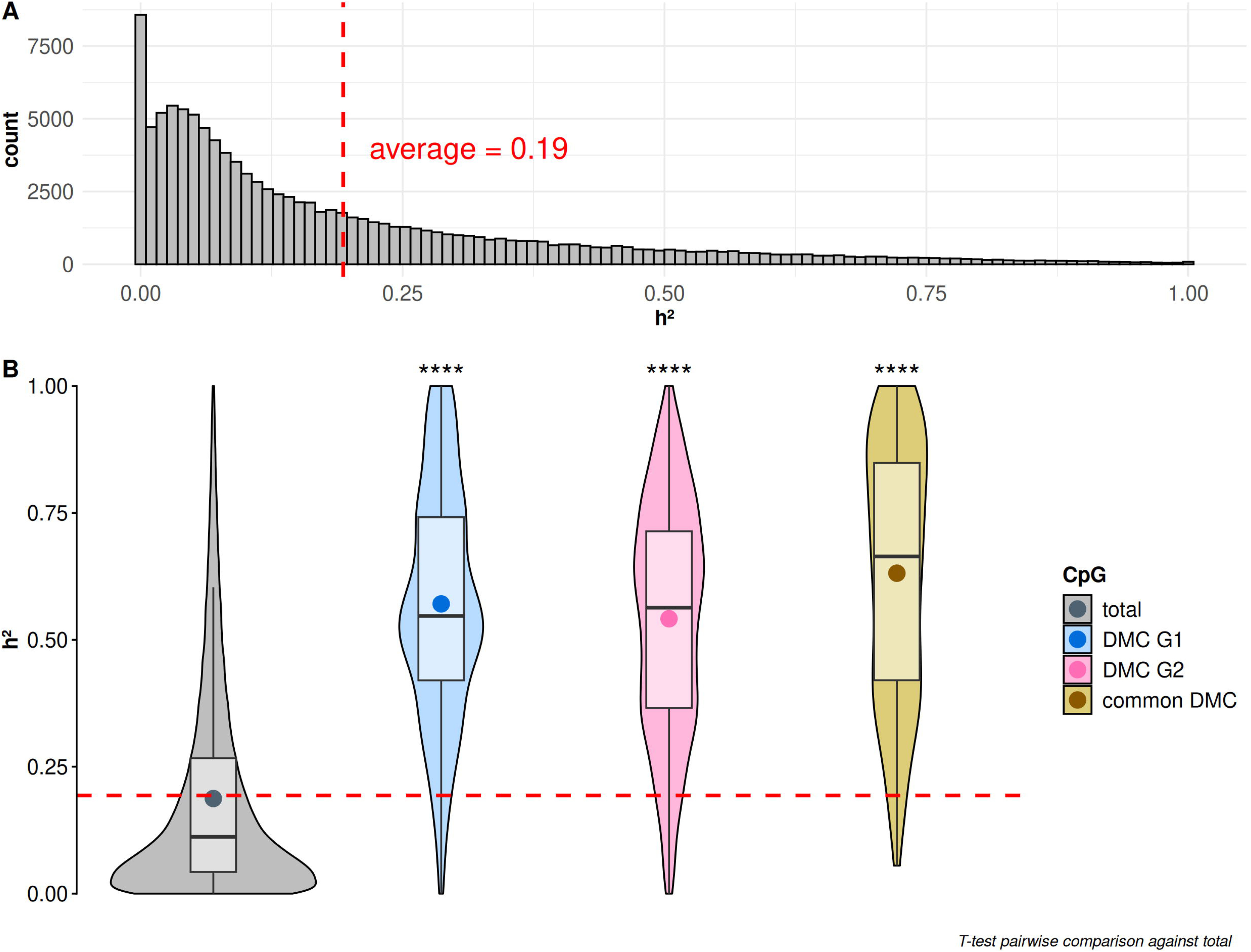
Heritability estimates of DNAm at CpG sites. A: Global distribution of heritability. The estimated average among all CpG sites is represented by the red dashed line. B: Pairwise comparison of heritability estimates at CpG sites identified as DMC in G1 (blue), G2 (red) or in common (beige). The T-test pairwise comparison was always done between the subset considered versus the totality of CpG sites (grey).

### Association between SNP and DNA methylation at DMC (meQTL)

To investigate the genetic determinism behind the regulation of DNAm at DMCs, meQTL analyses were performed for common DMCs in G1 and G2 (Figure 7). Out of the 133 DMCs identified on annotated chromosomes, 123 were found to be associated with at least one SNP. On average, DNAm at DMC was associated in cis with 47.3 SNPs within 1Mb, associated in trans with 0.5 SNPs located on the same chromosome (> 1Mb) and associated in trans with 1.3 SNPs located on another chromosome. These results suggest a strong genetic basis of the regulation of DNAm at DMCs. Ten DMCs were without cis and trans association with the SNPs available in our RRBS data set. They appeared to be located in regions where SNP coverage was limited. Combining both the NCBI and Ensembl annotations for the reference genome, they were located on chromosome 13 inside a gene, LOC107320288, annotated as protocadherin beta-15-like, and a long non-coding RNA (ENSCJPG00005011489), and on chromosome 21 in the *PRDM16* gene (PR domain containing 16).

**Figure 7:**
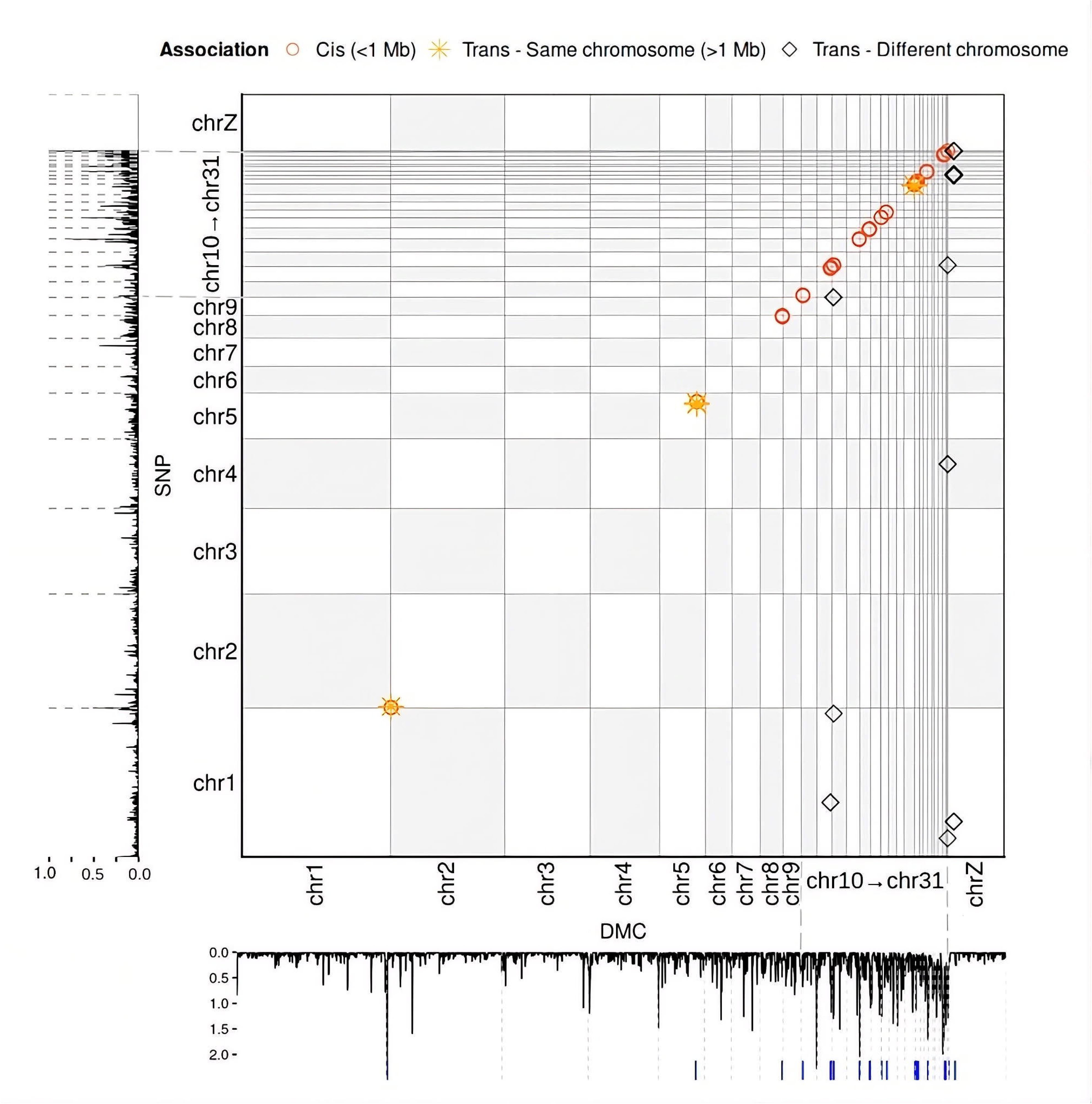
meQTL results. Association between DNAm at DMC (columns) and SNPs (rows). Only the significant associations were represented. The margin plots represent the CpG sites density along the genome (bottom graph) with the blue dashes representing the position of DMCs. On the left side, SNP positions along the chromosomes are represented. Only SNPs from autosomes were considered in the analysis.

## Discussion

The present study analysed the DNA methylation profiles from 1,261 Japanese quails of the same experimental design across five successive generations. Quail is a highly valuable model species for studying changes across generations, as its short generation interval and easy rearing, in terms of space and resources, make it particularly suitable for multigenerational experiments, as reflected by its use in numerous previous studies.(46,56–59). Here, we use a large animal design to obtain extensive data across several generations for genetic and epigenetic analysis.

### Global methylation landscape in red blood cells of Japanese quails

DNA methylation results were obtained using reduced representation bisulfite sequencing (RRBS) on red blood cell samples. RRBS is a widely used method for DNA methylation analysis that covers around 2 to 5% of the genome, in regions enriched in CpG sites and regulatory regions (60). Consistent with expectation, the distribution of the analysed CpG sites across genomic features showed enrichment in promoters, 3’UTR regions, coding sequences and intergenic regions directly located downstream of the CDS (< 1kb), thereby increasing the representation of genomic regions with potential functional relevance.

Globally, the mean methylation rate at CpG sites was low, with the majority of sites exhibiting little to no variation. This distribution differs from classical patterns observed in a wide range of tissues in mammals, where CpG sites generally display a methylation rate distributed in a U-shaped curve (11), but has previously been reported in avian species (61), including quail (46).

DNAm rate depended on the genomic feature, regardless of the genistein ancestral exposure. Such an observation was expected, given the strong dependence of gene regulation and expression based on methylated states of CpG sites. Unmethylated promoters are usually associated with a transcriptional active state, while methylated gene bodies can enhance transcription activity (62,63), explaining the patterns found of DNAm rates across genomic features in this study.

### Biological relevance of low methylation differences between epilines

The methylation differential analysis revealed DMCs and DMRs in generations G1 and G2 between epilines. The large sample sizes available within each epiline provided sufficient statistical power to detect significant differences even when the magnitude of the methylation difference was modest. Indeed, some of the CpG sites that were significant DMCs and DMRs showed very small differences in mean methylation between the epi− and epi+ lines, sometimes around 1%. To our knowledge, this quail design is among the largest designs published to perform differential analysis. Other DMCs displayed larger differences in mean methylation rates, up to 20%. Although the biological relevance of such differences remains to be established, small DNAm differences have already been reported in association with population differentiation (64), or with different phenotypic outcomes associated with lifestyle (65,66). In cancer, small DNAm differences can have drastic consequences, depending on the genomic context. Therefore, the magnitude of a methylation difference alone may not necessarily reflect its biological relevance. This may be particularly important when differences occur within regulatory regions. Thus even relatively small differences in DNA methylation could potentially contribute to phenotypic variation.

Seven genes were shared among generations and associated with DMC and DMR (four in gene body, three in promoter). They span functionally unrelated processes rather than a single pathway, based on information from GEGA (67) and GeneCards (68). This is consistent with the observed absence of significant GO enrichment. In the gene body, *DHPS* (Deoxyhypusine Synthase) regulates protein maturation and translation and is involved in the regulation of cell population proliferation, *GCDH* (Glutaryl-CoA Dehydrogenase) is involved in fatty acid beta-oxidation, *JPH3* (Junctophilin 3) maintains endoplasmic reticulum/plasma membrane junctions in neurons and is involved in neuromuscular processes, and *WDR83* (WD Repeat Domain 83) acts as a scaffold protein organizing multiprotein signaling complexes. In the promoter, *ARHGDIG* (Rho GDP dissociation inhibitor gamma) regulates Rho-GTPase activity with roles in cell adhesion and protein localisation, *NAA80* (N-Alpha-Acetyltransferase 80, NatH Catalytic Subunit) mediates N-terminal acetylation of actin, and *NUDT16L1* (Nudix Hydrolase 16 Like 1) modulates the DNA damage response by the negative regulation of double-strand break repair. Overall, these genes do not appear to reflect a coordinated biological programme.

However, the annotation of DMCs and DMRs common to G1 and G2 revealed several genes with potential relevant biological functions, providing candidate loci with persistent DNA methylation differences following genistein supplementation.

Among the set of gene-body DMC positions shared across generations, *NOTCH1* (Notch Receptor 1, a cell-fate receptor involved in developmental processes) can be related to genistein: the expression of this gene has been shown to be inhibited by genistein in a colon cancer model (69). For *WLS* (Wnt Ligand Secretion Mediator, essential for Wnt signaling) the two-generation stability was observed at the DMC level, and a DMR signal was also detected in generation 2. Wnt signaling can be modified by genistein during colon cancer development, notably through epigenetic modulations, including DNA methylation (70,71). *AR* (androgen receptor), which encodes a transcription factor mediating androgen-responsive gene expression, was differentially methylated in generation 2. It belongs to the steroid-signalling axis, for which a promoter-methylation silencing mechanism is well documented (72). Interestingly, genistein has also been reported to act as a tissue-specific androgen receptor modulator (73), mediated by *ESR2* (74). Genistein increases estrogen receptor beta (ER-β) levels via reducing its promoter methylation and ER-β, in turn, mediates the preventive action of genistein (75).

*HYAL3* (Hyaluronidase 3), differentially methylated at the promoter in both generations at both single-CpG and regional resolution, encodes a hyaluronidase involved reproductive functions, as sperm function and follicular atresia, and fertilization. Interestingly, a modest epiline effect on reproductive precocity, measured as age at first egg, was previously reported in the same experimental population (41,76). Given the role of phytoestrogens in regulating female reproductive function, *HYAL3* may represent a candidate mediator of genistein-induced epigenetic effects on reproductive pathways, although this hypothesis remains to be functionally tested (77).

Finally, *PRDM16* (PR/SET Domain 16) encodes a transcriptional regulator with histone methyltransferase activity and could be considered as shared across generations, as DMCs fell close to (Ensembl) or within (NCBI, data not shown) this gene. It encodes a chromatin-associated transcriptional regulator that governs brown/beige adipocyte identity and thermogenic gene expression through direct recruitment of histone- and DNA-modifying complexes in mammals (78) and preadipocyte proliferation in chicken (79). Together with *MGRN1* (Mahogunin Ring Finger 1, an E3 ubiquitin ligase), another gene acting on melanocortin receptor signalling and appetite regulation, *PRDM16* could be a plausible molecular candidate for the progressive divergence in body weights observed between epilines (79,80).

When broadening the analysis to include all DMCs for each generation, additional interesting elements emerge. Genes involved in embryonic developmental signalling pathways are found (*NOTCH1*, *WLS*, *BMP4…*). Development, differentiation and their regulation appear to be particularly affected in G2, as DMC or DMR are located within genes specific to these functions (*BMP4*, *FOXE3*, *FOXP1*, *GSX2*, *LBX2*). This may reflect a recalibration of early cell-fate decision networks, potentially underlying variation in growth, reproduction or behaviour, as previously observed in other quails’ phenotypes (56) including in the present experimental design (41). Genes encoding signalling components and transcription factors (*NF2*, *ERBB4*, *SOCS1*, and *OLIG3*, *TBR1*, *MYCL*, *SKOR1*) were also represented in G2. A substantial number of genes associated with the cytoskeleton, motility and cell adhesion were found in both G1 and G2 (*RHOA*, *RAB11FIP3*, *DNAI1*, *ANO8*), although some were Overall, the functional annotation of the DMCs and DMRs shows that the epiggeneration-specific (*MAST3* in G1; *CIMIP3* or *PRKCA* in G2). Several of these genes relate to ciliary function, which is consistent with the presence of genes linked to reproduction and gonadal function, such as *HYAL3*, *AR* or *DNTTIP1*. Sperm analysis in our lines could be a valuable follow-up to test for an effect on male reproduction. Finally, given that genistein is itself a methylation modifier, it is worth noting that genes involved in DNA methylation or chromatin conformation showed differential methylation between epilines, most of them in G2 (*DNMT1*, *TET2*, *PRMT5*, *INO80B*), with the exception of *MBD2*, which is G1-specific. That these epigenetic regulators are themselves differentially methylated suggests a possible feedback loop: the initial exposure may have altered the expression of proteins that, in turn, shape the methylation landscape transmitted to subsequent generations.

This functional annotation of DMCs and DMRs shows that the epigenetic signature associated with maternal genistein supplementation is not organised around a single, statistically dominant functional module. This result must be interpreted in light of two technical limitations: the relatively small size of the gene lists considered, which limits the statistical power of this type of test, and the ascertainment bias inherent to RRBS, which preferentially targets CpG-rich regions already known to be associated with housekeeping and developmental genes (81,82). Despite these limitations, the few candidates found could support DNAm as a potential candidate mechanism for transmitting environmental effects, especially genistein-derived, in subsequent generations.

### DNAm as a candidate mechanism for environmentally induced epigenetic inheritance

The main objectives of this study were to (1) verify the existence of differential DNAm profiles between control and genistein-derived quails and (2) assess their evolution across generations. Surprisingly, no differentially methylated cytosines (DMCs) or regions (DMRs) were identified in G0, the first generation after the environmental perturbation. By contrast, in G1, 621 DMCs were identified (located in 36 DMRs) and this number further increased to 1381 DMCs (located in 101 DMRs) in G2, among which only 166 DMCs (13 DMRs) were in common. This progressive increase and molecular differences between epilines interestingly echoes other observations from environmentally-induced multigenerational epigenetic models, as well as previous findings from the same experimental population, in which differences in body weight between epilines increased across generations (41). In rats exposed to vinclozolin, DMRs identified in the sperm of directly exposed individuals were fewer and mostly distinct from the DMRs identified in the sperm of their progeny, two generations later (24). For this particular example, the authors discuss the possible impact of vinclozolin in altering the developmental programming of germ cells in the progeny, hence resulting in different yet accumulating molecular differences. We hypothesise that the mechanism involved could be a so-called ‘wash-in’, meaning that methylation slowly accumulates over successive generations, as observed in other species (83). In another study also in rats, environnementally-induced epigenetic modifications favoured the appearance of copy number variants in later generations, hence resulting in a transgenerational phenotype of both epigenetic and genetic basis (84). Indeed, the increase of differences in DNAm profiles observed between epilines across generations could originate from an ancestral genistein-induced molecular change transmitted to the progeny, genetic differences between epilines influencing DNA methylation patterns, or a combination of both epigenetic and genetic mechanisms of inheritance. The use of fixation index (F_ST_) to compare epilines based on bi-allelic differences provided some insight on that matter. Mean F_ST_ between epilines and generations were low (0.004 and 0.003 respectively, Supplementary Table5) showing little to no overall differentiation, inferior and much lower compared to values known for intra-breed differentiation in livestock (85). Additionally, moderate signatures (< 0.2) between G0 and G2 individuals, on chromosome 2, 4 and 15, regardless of their epiline were identified, while no specific signature between epilines was detected. Although F_ST_ only provides a relative measure of population differentiation, whose magnitude depends on the distribution of allele frequency and within-population diversity (86), the low F_ST_ observed between epilines reflect that the mirror-mating design was effective in limiting genetic drift and maintaining genetic similarity between the two epilines across generations. Interestingly, while DNAm differences between epilines increased across generations, this was not accompanied by a corresponding increase in genetic differentiation. This observation argues against genetic factors as the sole contributors to explain the increasing DNAm differences across generations, although genetic effects at specific loci cannot be excluded. These results are therefore consistent with DNAm being a plausible molecular candidate involved in the transmission of environmental effects across generations. These findings further question the extent to which DNAm differences are under genetic control.

### Genetic determinism of DNA methylation

SNP-based heritability for DNA methylation was estimated for each CpG site, and averaged 0.19, suggesting that genetic factors contribute to variation in DNA methylation. To our knowledge, these represent the first SNP-based heritability estimations of DNA methylation in an avian species. These results were consistent with heritability estimations of DNAm in human blood (34,87), as well as in other studies in livestock, in different tissues (88,89). Although no avian study has directly reported whole-genome heritability estimates for DNAm, works evaluating the contribution of individual SNPs to variation in DNAm levels have found genetic effects of similar magnitude (28,84,85). However, our estimates were obtained from CpG sites captured by RRBS, thus biased towards a fraction of the genome enriched in regulatory regions. Interestingly, the mean heritability estimates for DNAm at CpG sites found significant in the differential analysis were significantly higher than the overall mean estimations. Similarly, DNAm in functional regions in cattle was slightly more heritable (0.36) than previously reported genome-wide (90). The distribution of DMCs in genomic features could explain these higher values, given the strong proportion of DMCs located in promoters (Supplementary Figure6). Indeed, we hypothesise that changes in methylation rate at CpG sites in promoters could impact the regulation of associated gene expression.

The meQTL analysis of DMCs provided further information on the genetic determinism of DNAm variation. Because SNPs were called from the RRBS data, genome SNP coverage was more limited compared to whole-genome SNP datasets. This limited SNP coverage reduces the ability to detect meQTL associations and may particularly affect the detection of trans-meQTLs, for which adequate genome-wide SNP coverage is important. The absence of recorded associations for 10 DMCs could therefore, at least partly, reflect insufficient SNP coverage rather than the absence of genetic effect on DNAm variation at these sites. SNP-calling from bisulfite sequencing remains technically challenging, as it requires distinguishing genuine C/T polymorphisms from C/T conversions induced by bisulfite treatment. BISCUIT is a recent tool (45) among the few that were specifically developed for SNP-calling from bisulfite data (91,92) and has been shown to achieve high accuracy in SNP calling from RRBS data (93), particularly in non-model organisms (94).

These findings suggest that DNAm at differentially methylated CpGs between epilines is often associated with close-by genetic variability (cis-association) across multiple SNPs. Overall, our meQTL results at common DMCs aligned with previous studies, as methylation was usually under the regulation of several SNPs. The underlying genetic regulation of DNAm was indeed previously reported in other avian meQTL studies (28,95), where authors propose the implication of DNAm in adaptation and domestication being most likely linked to underlying genetic variations. In the present study, we globally demonstrated that DNAm at CpG sites showing differential methylation were extensively under genetic control.

### How this study opens up new avenues for the analysis of multigenerational genetic and epigenetic

To conclude, this work revealed differentially methylated cytosines (DMCs) and regions (DMRs) among two groups of Japanese quails with different ancestral environmental backgrounds. Some of the significant DMCs detected were surprisingly associated with low differences in mean methylation levels between groups, raising questions about their biological relevance. Strikingly, a subset of DMCs persisted across generations and these profiles were largely controlled by an underlying genetic background. However, no apparent genetic differentiation between epilines was captured in our data, uncovering a genetic-epigenetic paradox. The increase in differences between epilines across generations possesses a genetic basis with too small effects to distinguish epilines, yet sufficient to potentially induce different molecular outcomes. Besides, if DNAm at CpG sites were totally controlled by genetic factors, similar observations would also be expected in generation G0. This increased differentiation between epilines throughout generations requires further investigation. Recent findings on epigenetic reprogramming in birds seem to demonstrate the absence of global erasure of methylation profiles across embryonic stages (96). Under the light of our results, we could hypothesise that DNAm profiles are partly conserved from one generation to the next, through genetic or non-genetic transmission of the methylation information, and that such positions undergo an accumulation of methylation as previously proposed (24,84).

An important next step will be to determine whether these differential methylation patterns influence gene regulation and ultimately contribute to phenotypic variation. Integrating complementary transcriptomic data with genotypes, DNA methylation and phenotypic records would help to understand the functional consequences of the observed methylation differentiation and assess whether relatively small differences in methylation levels may have phenotypic outcomes, especially as increasing phenotypic differences have already been reported in this current dataset (41). More broadly, the combination of molecular and phenotypic information available in this large multigenerational Japanese quail design provides a unique framework to study the interplay between genetic and epigenetic mechanisms in shaping phenotypic variability, and their transmission throughout successive generations.

## Acknowledgements

The authors would like to express their sincere appreciation to Marie Courbariaux, Florence Jaffrezic, Andrea Rau and Guillaume Cossard for their expertise on methylation differential analysis. The authors also address special thanks to Alexandre Hubert, Pedro Sà and Maria Luigi for their precious contributions and proficiency during the numerous DNA methylation analysis workshops that were organised throughout this study. Last but not least, many thanks to the animal caretakers at UE1295 PEAT (Nouzilly, France, https://doi.org/10.15454/1.5572326250887292E12) for their commitment to the quails’ welfare, as well as the Genobioinfo platform for the access to their computation environments, maintenance of the data center and their technical support.

## Disclosure of interest

The authors report there are no competing interests to declare.

## Declaration of generative AI use

The authors did not resort to using AI or AI-assisted technologies for this work.

## Funding details

This project has received funding from the European Union’s Horizon 2020 research and innovation program under grant agreement N°101000236 (GEroNIMO). This project is part of EuroFAANG (https://eurofaang.eu). Stacy Rousse is co-funded by the INRAE Animal Genetics Division and the French Occitanie Region. Stacy Rousse received financial support from Toulouse INP-ETI (EpiMob) for a four months research stay at Wageningen University & Research.

## Authors contribution

SR actively took part in the conceptualization, methodology, investigation and visualization, as well as performed data curation and conducted most formal analyses. SLe contributed to the investigation and provided resources by creating all RRBS libraries. RS participated in the data curation and management of resources through the software development of the RRBS nextflow pipeline. DG was in charge of the animal facility and provided resources for this research. MG and OM welcomed SR in their lab and participated in elaborating the research methodology, as well as took part in the investigation and validation of this work. SLa and TZ contributed to the validation and project administration as well as to manuscript revision, FP and SE supervised and contributed to the conceptualization, methodology, investigation and conducted formal analyses, as well as secured funding. SR, FP and SE did the writing of the original version, review and editing of the manuscript.

## Data availability statement

All raw data are available on Zenodo (doi: 10.5281/zenodo.22690759). The fastq files are available at ENA (https://www.ebi.ac.uk/ena/browser/home) with accession number PRJEB124227. All codes to perform analyses are available under the git repository: https://forge.inrae.fr/stacy.rousse/dnamethylation_inheritance_quails/

## Supplemental material

Supplementary Figure1: Summary of filters applied to RRBS data. A: Diagram summarising the filters and sample sizes for SNP and CpG filtering steps. Red: SNP data; Green: CpG data. B: Filters applied to CpG sites and comparison to total number of CpGs on the quail genome. From left to right: Total number of CpG sites annotated on the quail genome; number of unique CpG sites retrieved in the RRBS experiment; number of CpG sites common to 80% of samples; number of sites remaining after removing CpGs overlapping with SNPs; final number of CpGs after keeping sites with a standard deviation of the methylation rate ≥ 0.05.

Supplementary Figure2: Mean methylation differences between epilines at common DMCs and evolution from G1 to G2. The dashed line is y=x, the solid line is the regression line from the data. Globally, we see a conservation from G1 to G2 of the methylation profile at common DMCs (hypomethylated CpGs in G1 remain hypomethylated in G2, respectively hypermethylated). The slope indicates a slight tendency towards the increase of mean methylation differences, in either group, across common DMCs.

Supplementary Table1: DMR details. Columns are DMR: DMR identifier; chr: chromosome; start: start position of DMR; end: end position of DMR; size (bp): DMR size in base pairs; generation: generation in which the DMR is detected; nCpG: number of CpG contained in the DMR; nDMC: number of DMC contained in the DMR

Supplementary Figure3: Localization of DMCs detected for the sex covariate. A substantial number of sex DMCs were identified in all generations. An important proportion of identified DMCs were localized on the sexual chromosome Z (pink) in all generations. This distribution was expected, and supported the validity of the differential analysis results.

Supplementary Figure4: Permutation analysis for DMC detection: The distribution of the number of DMCs detected with simulations through permutation analysis (light blue) versus real observation of detected DMCs (red dashed line). The empirical probability to the risk alpha=0.05 statistically tests the potential stochastic detection of DMCs through the assumption H0: the number of observed DMCs comes from a similar distribution as the distribution of DMCs obtained with random permutations. p=r+1/n+1 where n is the total number of simulations and r is the number of simulations that detect a similar or higher number of DMCs than observed without permutation. All the DMCs detected in real observed data were significantly different from stochastically detected DMCs in permutations, except for the epiline in G0, because no DMC for the epiline was observed in this generation. For that reason, all simulations exhibited an equal or higher number of detected DMCs, resulting in a pvalue of 1.

Supplementary Table2: NCBI list of genes associated with DMCs/DMRs. Supplementary Table3: NCBI list of promoters associated with DMCs/DMRs. Supplementary Table4: Ensembl list of genes associated with DMCs/DMRs. Supplementary Table5: Ensembl list of promoters associated with DMCs/DMR

Supplementary Figure5: Genome-wide F_ST_ values in RRBS SNPs. The population subsets of G0 versus G2 individuals and epi− versus epi+ individuals (lower tracks) were completed with F_ST_ computations between generations per epiline to check if any change in bi-allelic differences between generations aroused in a particular epiline (upper tracks). Thus, F_ST_ for epi− individuals between G0 and G2 and F_ST_ for epi+ individuals between G0 and G2 was computed. The signatures found in the global G0-G2 comparison were also present when comparing G0 individuals with G2 individuals per group, suggesting an accumulation of allelic differences attributable to time. F_ST_ was computed per SNP, in all autosomes (chr1-chr28). The two strips before chrZ correspond to the linkage groups chrLGE22C19W28_E50C23 and chrLGE64 respectively.

Supplementary Table6: Mean and weighted F_ST_ values for bi-allelic comparison of population subsets (epi− versus epi+, G0 versus G2, epi+ individuals in G0 versus epi+ individuals in G2 and epi− individuals in G0 versus epi− individuals in G2).

Supplementary Figure6: Repartition in genomic features of common DMCs identified in G1 and G2.

## Notes

### Competing Interest Statement

The authors have declared no competing interest.

