## Supplementary figures and images for "Genetic determinism and inheritance of differential DNA methylation profiles across three successive generations in quail"

### Supplementary Figure1

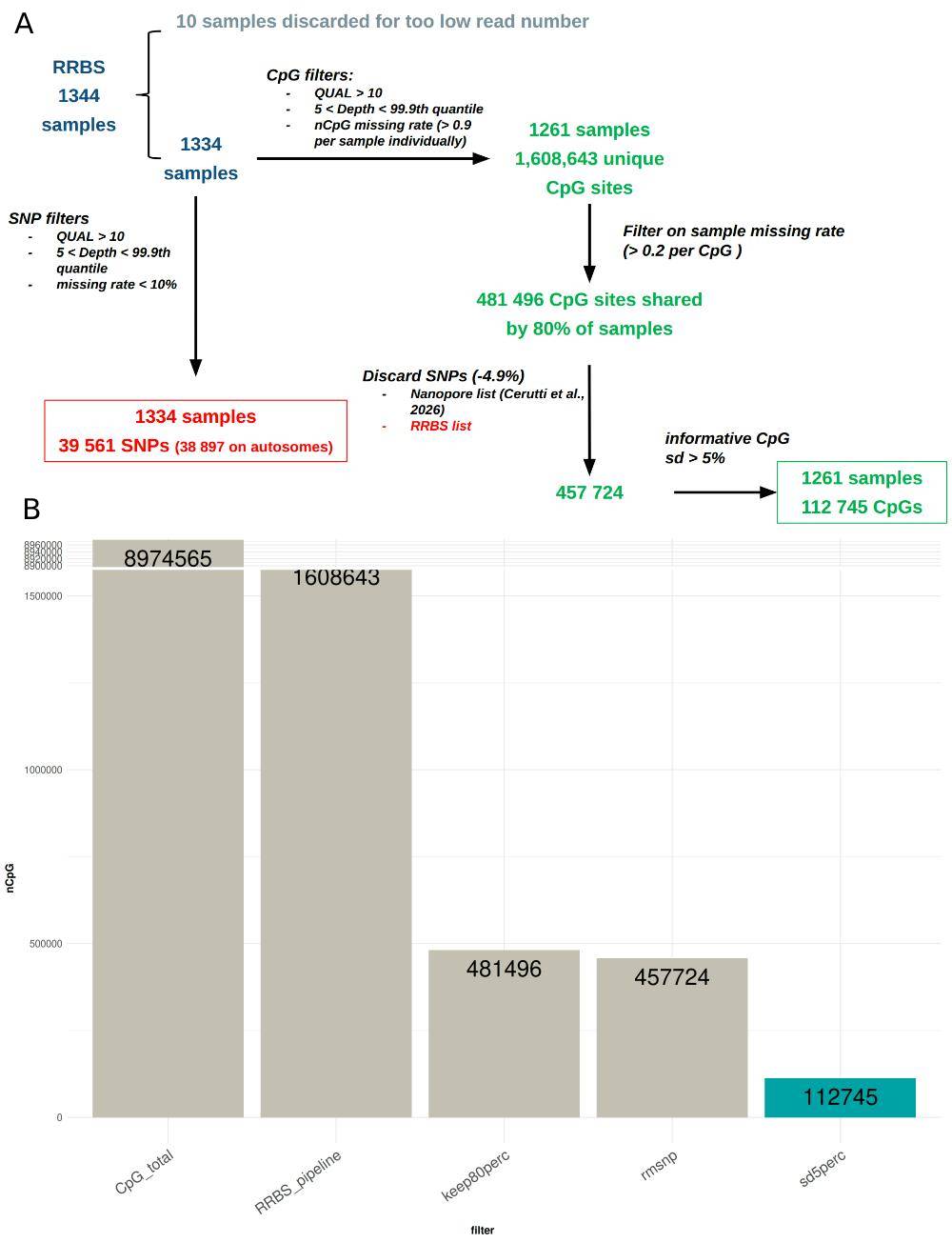

### Supplementary Figure2

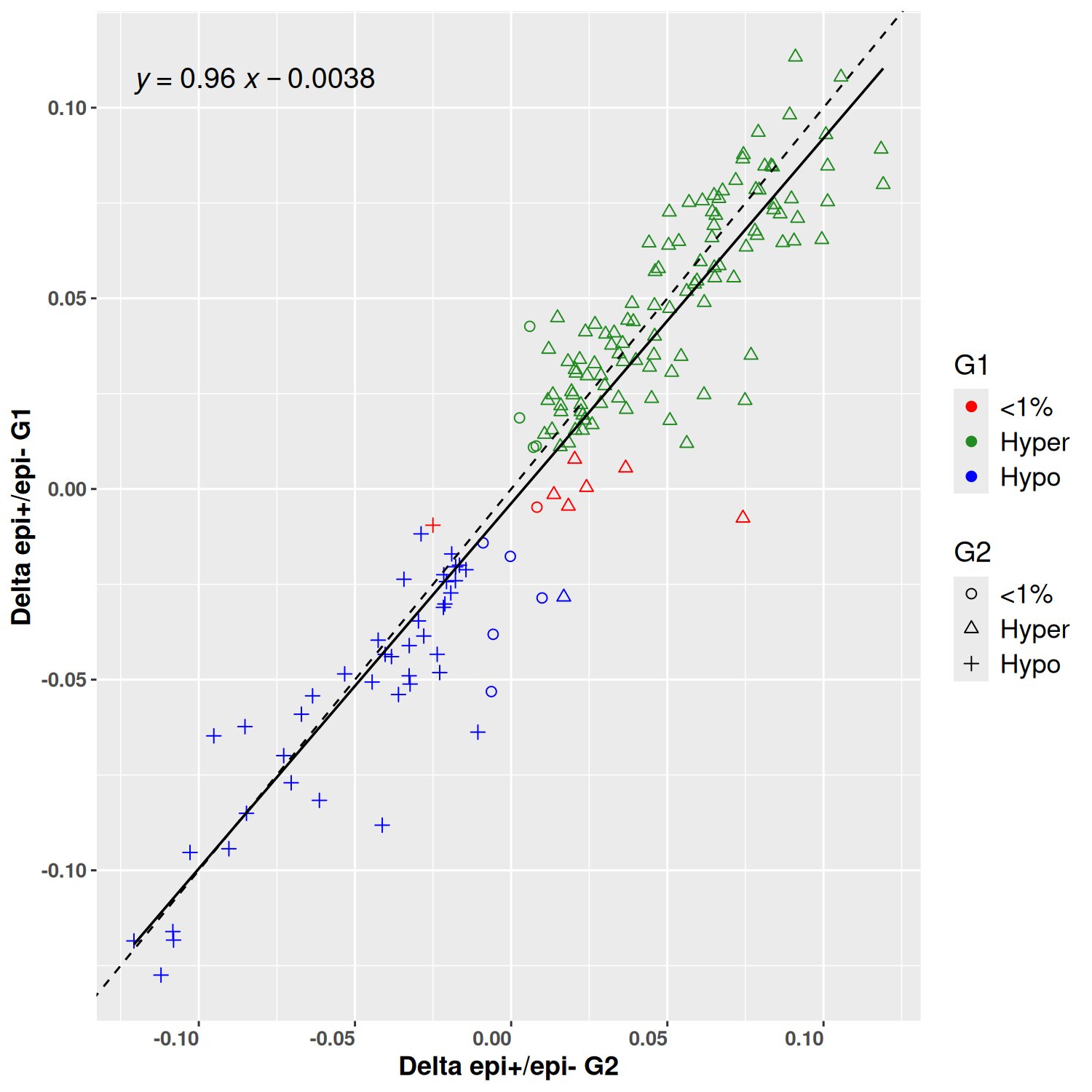

### Supplementary Figure3

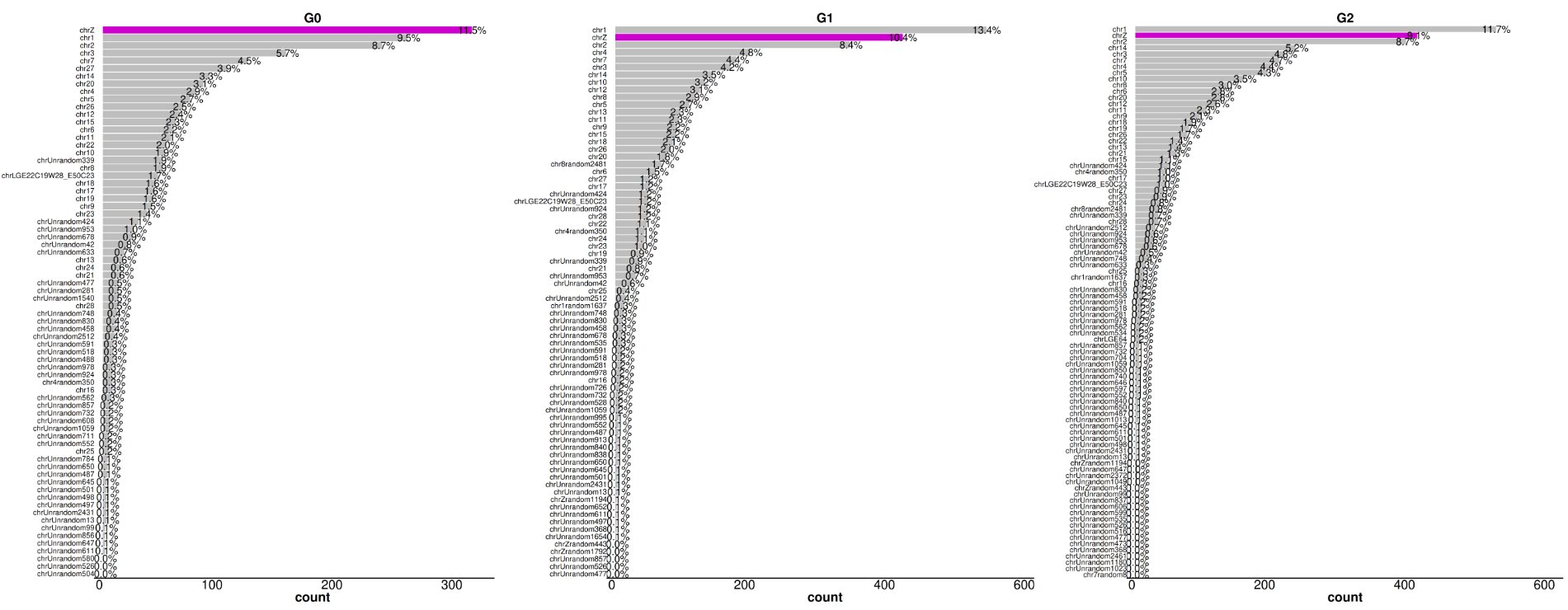

### Supplementary Figure4

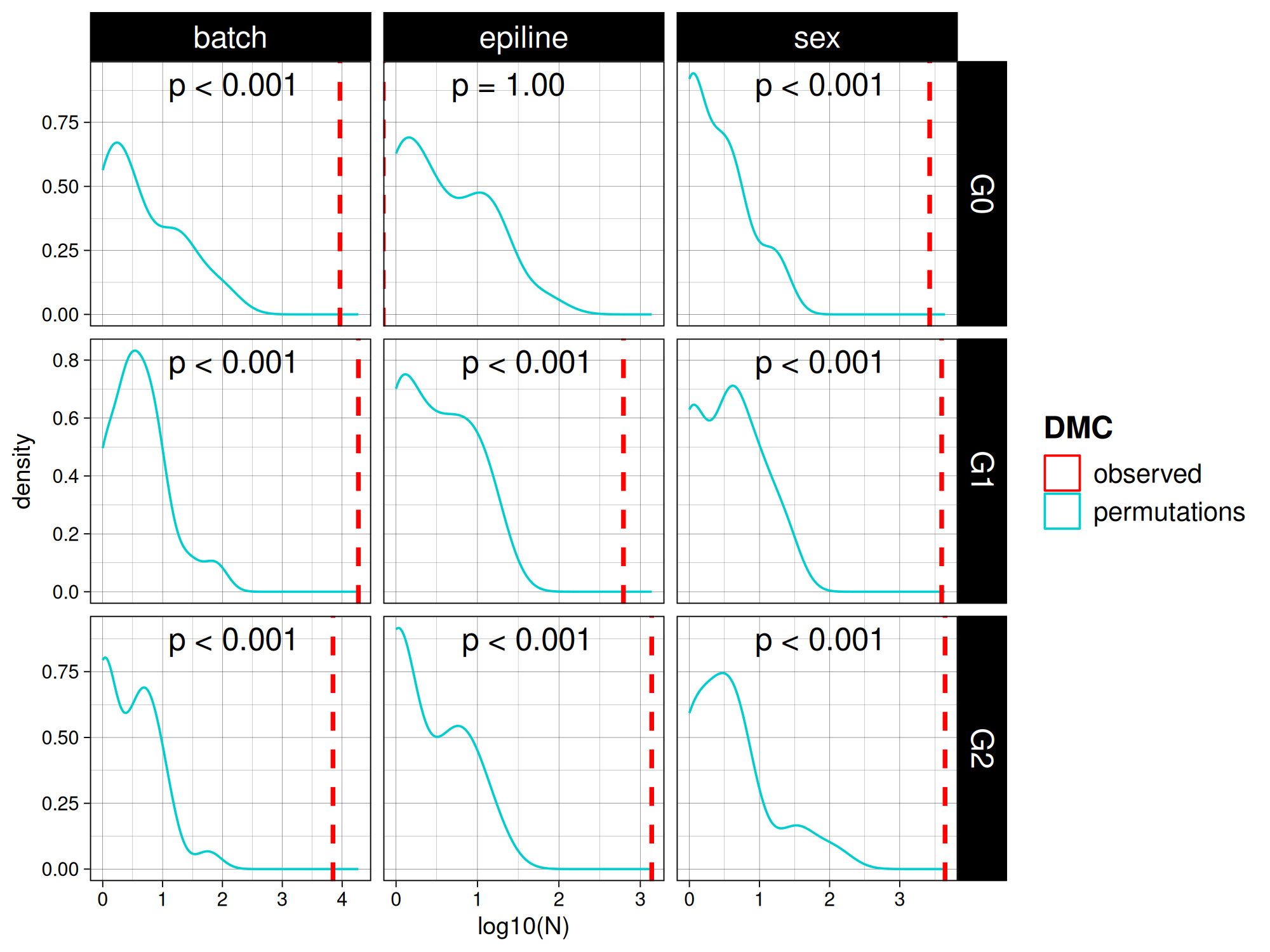

### Supplementary Figure5

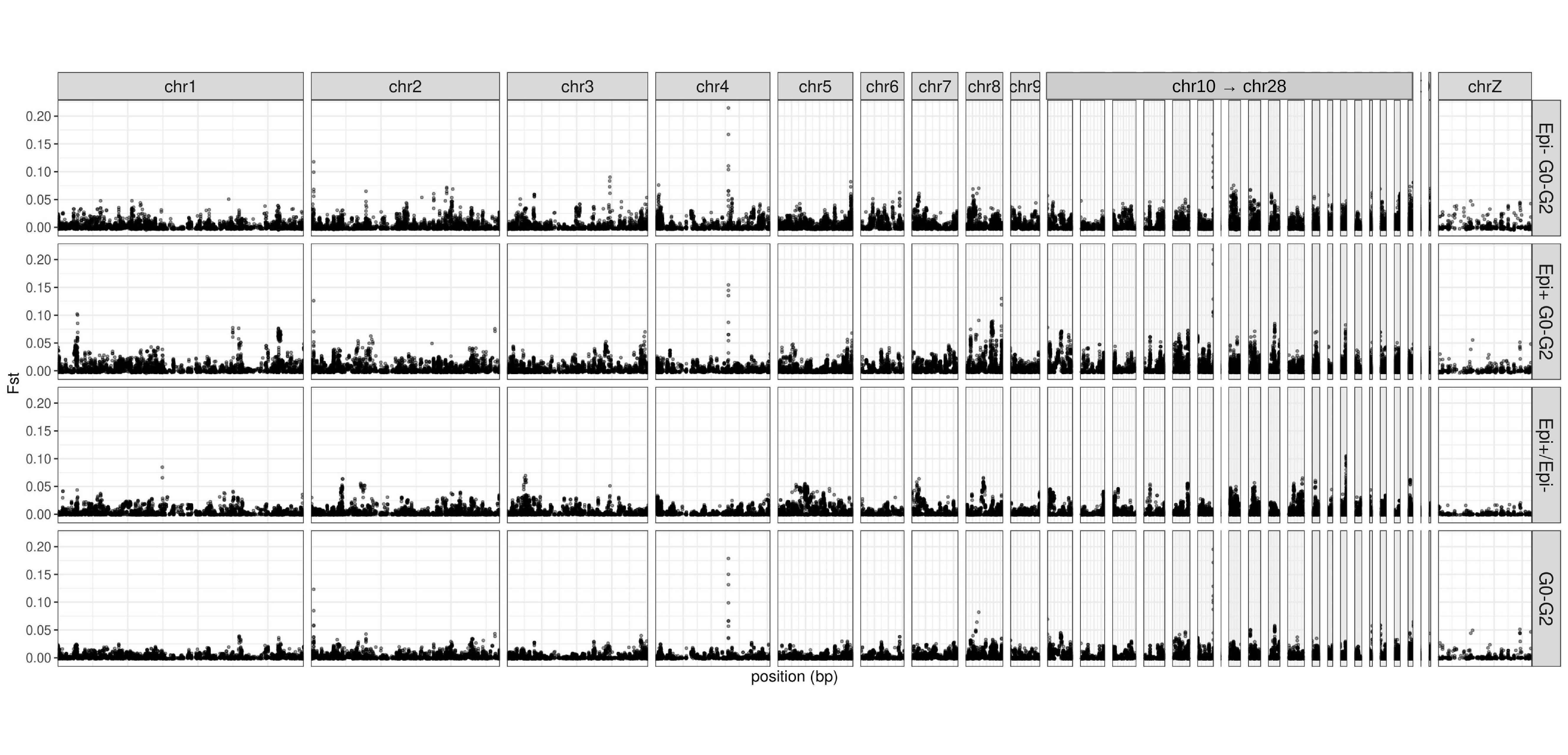

### Supplementary Figure6

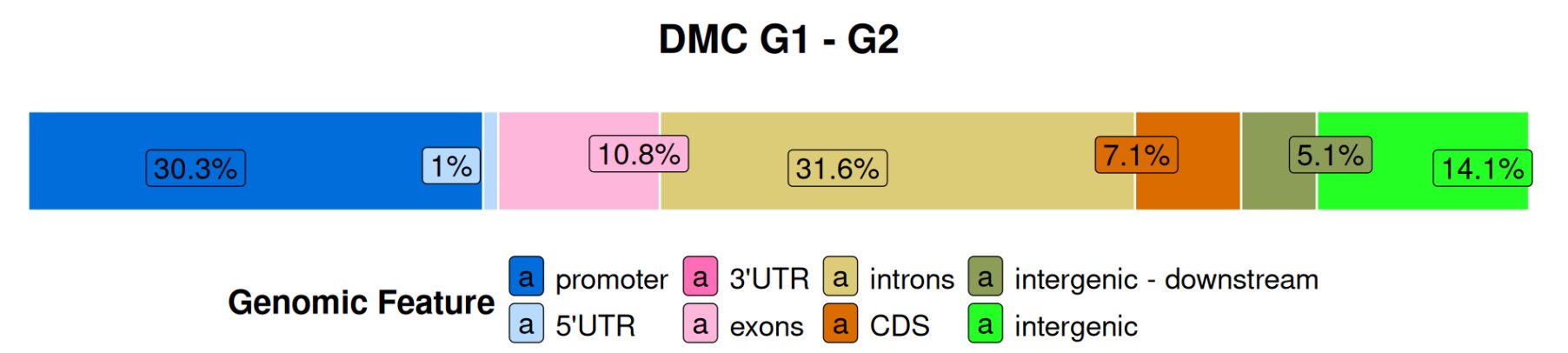
